# Urequinona, a molecule from the root of *Pentalinon andrieuxii* Muell-Arg heals *Leishmania mexicana* ear’s infection in mice: This plant is widely used by the Mayan traditional medicine

**DOI:** 10.1101/2021.03.02.433683

**Authors:** AP Isaac-Márquez, CM Lezama-Dávila

## Abstract

In this work we tested both the *in vitro* and *in vivo* anti-*Leishmania mexicana* activity of a molecule we originally identified in the root of *Pentalinon andrieuxii* Muell-Arg, a plant that is widely used in Mayan traditional medicine. The chemical name of this molecule is 24-methylcholesta-4-24(28)-dien-3-one and for simplicity we assigned the short and trivial name of urequinona that will be used throughout this work. It induces necrosis and apoptosis of promastigotes cultured *in vitro* and extensive ultrastructural damage of both promastigotes and amastigotes. It also induces production of Interleukin (IL)-2 and interferon (IFN)-γ by splenic cells from infected and urequinona treated mice stimulated *in vitro* with parasite antigen (Ag) but inhibits production of IL-6 and IL-12p70 by bone marrow derived macrophages (BMM) infected *in vitro* and then treated with urequinona. It also induces activation of transcription factors such as NFkB and AP-1 (NFkB/AP-1) in RAW reporter cells. We also developed a novel pharmaceutical preparation of urequinona encapsulated in hydroxyethyl cellulose for dermal application that significantly reduced experimentally induced ear′s lesions of C57BL/6 mice. We conclude the preparation containing this molecule is a good candidate for a novel anti-leishmanial drug′s preparation.

## Introduction

Leishmaniasis is a parasitic disease provoked by different species of the genus *Leishmania* (1,2). *Leishmania mexicana* (*L. mexicana*) is responsible for a highly endemic clinical condition found in Central America, Mexico and some parts of Southern United States (3,4). In the Yucatan peninsula this condition is also known as chiclero’s ulcer that evolves with a single cutaneous lesion at the site of an infected sanfly bite and is medically described as Localized Cutaneous Leishmaniasis (LCL,3,4). Antimonial drugs are the first line drugs prescribed to treat this medical condition but other ones currently in use include amphotericin B, pentamidine, paromomycine (5-8). However, all these compounds present high level of toxicity and, since intramuscular injections of antimonial drugs are painful, patients frequently abandon their treatment that, on the other hand, is frequently un-available in endemic areas of Mexico. In the state of Campeche in Mexico, patients frequently rely on Mayan traditional medicine (9,10). We have tested a plant that traditional healers prescribe to treat several medical conditions including LCL. It is known with the vernacular name of “contra-hierba” and with the scientific name of *Pentalinon andrieuxii* Muell-Arg (9). We have previously demonstrated that the hexanic extract of the root of this plant kills promastigotes and amastigotes *in vitro* and heals ear infection of C57BL/6 mice (10,11). In this work, we present comprehensive *in vitro* and *in vivo* data of a known sterol we previously identified in the roots of this plant that we found it presents considerable anti-*L. mexicana* activity *in vitro* (10). We showed that this molecule we named it with the short and trivial name of urequinona heals ear induced infections of C57BL/6 mice. We also present data showing that this molecule provokes necrosis and apoptosis of *L. mexicana* promastigotes, induces extensive ultrastructural damage of promastigotes and amastigotes internalized by bone marrow derived macrophages (BMM) and activates transcription factors such as NFkB and AP-1. We also showed that spleen cells from infected and urequinona treated mice produce more interleukin (IL)-2 and interferon (IFN)-γ. However, *in vitro* infected and urequinona treated BMM produce less IL-6 and IL-12p70. Results of these findings are discussed.

## Material and Methods

### Preparation of a pharmaceutical formulation containing urequinona for topical treatment of ear’s *L. mexicana* infection of mice

Pure 24-methylcholesta-4-24(28)-dien-3-one (for simplicity we provisionally use the trivial name of “urequinona” throughout this work) was kindly provided by Drs. Pan and Abelhamid and it was prepared as we described it somewhere else (10). The chemical structure of this molecule is shown in Fig 1. We used this compound to prepare a topical formulation that is suitable for animal and human treatment of cutaneous infections with *L. mexicana* parasites. To prepare this formulation we used a known amount of urequinona mixed with a nonionic, water soluble polymer with pH=6.5 (commercially known as Natrosol gel™). Our preparation contains 5% 2,3-dihydroxypropyl (9Z)-9-octadecenoate (also known as monoolein that facilitates dermal absorption), 5% propylene glycol and 3% hydroxyethyl cellulose (HEC) in sterile distilled water. Control topical preparation included all above described components except urequinona as active ingredient (vehicle). Two weeks after infection, animal treatment with this topical preparation was performed as described below.

**Fig 1.**
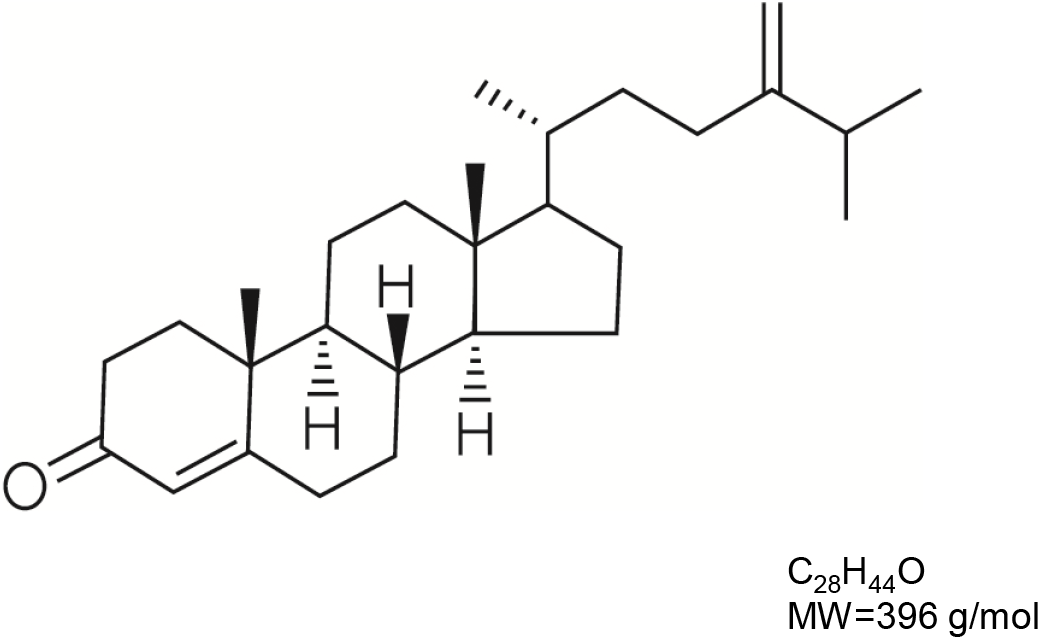
IUPAC name of tested compound: 24-methylcholesta-4-24(28)-dien-3-one. For simplicity we assigned it the short and trivial name of “urequinona”. Molecular formula and molecular weight (MW) and are also indicated on the right hand side of the image.

### Animals

C57BL/6 mice of both sexes (10-15 weeks old) were used throughout these experiments and were maintained and bred in an animal facility in accordance with bioethical guidelines. Institutional ethical approval was given to perform all animal experiments. Mice of both sexes were used for BMM isolation and *in vivo* infection using the ear model of infection (10,11).

### Parasites

A strain of *L. mexicana* parasites (WHO code MNYC/BZ/62/M379) was used in this work. We also used a line of transgenisc *L mexicana* (DsRed *L. mexicana*) expressing firefly luciferase and red fluorescent protein, originally provided by Prof. Beverly and prepared as follows. Clonal transfectants were developed by homologous integration of a LUC-DsRed2 cassette (SwaI fragment from plasmid pIRISAT-LUC-DsRed2 strainB 5947) into the ribosomal RNA (rRNA) locus as previously described (12). For simplicity we abbreviated these parasites as DsRed *L. mexicana*. Fluorescence intensity of transgenic parasites was verified by fluorescent microscopy in promastigotes and amastigotes (inside of BMM). All parasites (regular and transgenic) were cultured in complete media (RPMI-1640 containing 10% of fetal calf serum from Sigma, St. Louis, MO plus 100 µg/mL of erythromycin and 100 IU of penicillin from GIBCO BRL, Grand Island, NY). Parasites were kept by serial passage of amastigotes into the shaven rump of C57BL/6 mice. Highly infective metacyclic promastigotes were negatively selected by using the negative selection protocol of peanut agglutinin (PNA,10). In brief: 1.5 × 10^8^ promastigotes were treated with PNA for 30 min at room temperature and immediately after they were spun down at 200 X g for 5 min. Parasites from supernatant were collected, washed and counted followed by its suspension in saline solution and injected into the ear dermis of C57BL/6 mice. Two weeks after animal infection, topical treatment with our urequinona containing preparation was performed.

### Treatment of *L. mexicana* infected ears of C57BL/6 mice

Two weeks after initial infection, once lesions were visible and palpable, mice were treated topically every day for 12 weeks with the pharmaceutical formulation of urequinona containing 10 µM of this steroid per dose/day. Control animals were only treated with vehicle. Lesion growth was weekly monitored using a dial gauge micrometer (Mitutoyo 0-10 mm) for 12 weeks and parasite loads were calculated at the end of the experiment by direct counting of parasites recovered from infected ears and a representative ear sample, randomly chosen was sent to pathology for histological processing using the hematoxylin-eosin staining protocol (11,12).

### Evaluation of necrosis and apoptosis of promastigotes treated with urequinona

Promastigotes of *L. mexicana* were treated with urequinona and their killing was assessed by flow cytometry as we previously described it (10-12). Briefly, one million parasites suspended in 1 mL of complete media were seeded into 24 well-plates. Next, we added known concentrations of urequinona (0-100 µM) dissolved in 2% dimethyl sulfoxide (DMSO) in complete media from Sigma-Aldrich and parasite cultures were incubated at 28 °C for 48 h. Control parasite samples were only treated with 2% DMSO in complete media. At this point parasites were centrifuged at 2000 RPM for 3 min and the pellet was re-suspended in 100 µL of PBS containing 20 µg of propidium iodide or Anexin V (Sigma-Aldrich) and incubated for 1 h in the dark at 4 °C. The resulting parasite suspension was made up to 1 mL and 15,000 cells per sample were tested by flow cytometry using a FACSCalibur instrument from Becton Dickinson.

### Evaluation of killing of promastigotes and amastigotes treated with urequinona

Testing promastigote killing was performed by seeding 1 × 10^6^ parasites in 1 mL of complete media containing 10 µM of urequinona or sodium stibogluconate (reference drug) in 24 well plates (Corning Inc., NY). After 48 h of incubation, parasite killing was evaluated by flow cytometry as described above. In order to test intracellular killing of parasites we collected bone marrow from femurs and tibias of healthy C57BL/6 mice of both sexes. Marrow cells were disrupted aided by a sterile syringe plunger, pooled and cultured in a CO_2_ incubator (adjusted at 37°C and containing 95% air with 5% CO_2_) in complete culture media supplemented with 20% of supernatant from cultured L-929 cells. These cells were grown in 75-cm^2^ cell culture flasks (Corning Inc., NY) and collected after 5 days of culture. More than 90% of differentiated macrophages were positive for CD11b, a monocyte/macrophage marker as tested by flow cytometry (FACScalibur, Becton Dickinson, San Jose, CA, 11). Half million BMM were seeded on top of circular glass coverslips placed at the bottom (one coverslip/well) of 24-well tissue-culture plates (Corning, Inc.). Macrophages were infected overnight with 5 × 10^6^ promastigotes from stationary phase *L. mexicana* (ratio 10:1). Next, BMM were extensively washed with sterile PBS to eliminate non-internalized parasites. At this point infected macrophages were treated with different concentrations (0-100 µM) of urequinona or reference drug (0-400 µM) for a period of 48 h. After this time cells were stained with Giemsa, washed with PBS, fixed with methanol for 2 min. and mounted on a glass slide using a microscopy tissue mounting medium (Sigma-Aldrich) followed by microscopy observation using a 100X objective. We counted, in different microscopic fields, the number of parasites per 100 macrophages and the percentage of infected macrophages (10).

### Viability test of macrophages and splenocytes

Viability of both macrophages and splenocytes was carried out using the trypan blue test. Cells (10^5^/0.2 mL) were grown in 96 well plates inside of a CO_2_ incubator and were co-cultured with or without 100 µM of urequinona for 48 h. Next we spun down the cell suspension and pellet was re-constituted with 50 µL of complete media and mixed with 50 µL of trypan blue solution (Sigma-Aldrich). Plates were incubated at 37°C for 10 min to allow dead cells to internalize the stain. Finally, cell samples were observed under the microscope using the 40X objective. We counted two hundred cells per sample and recorded the number of dead cells with blue stained nucleus. Viability was expressed as the percentage of viable cells (without blue stained nucleus) and we found more than 90% of viability in all samples tested (10).

### NFkB/AP-1 reporter cell assay

A commercially available macrophage cell line known as Raw-Blue macrophages™ (InvivoGen Inc.) was employed in this experiment. These cells were transfected so they express a secreted embryonic alkaline phosphatase (SEAP) gene (stable transfectants) inducible by NFκB and AP-1 transcription factors as we previously describe it (12). Increase of enzymatic activity of SEAP is a direct indication of activation of these transcription factors. Therefore, we measured SEAP activity as follows: One hundred thousand cells in 200 µL of complete media containing 10 µM of urequinona or control solution containing 2% of DMSO were seeded into 96 culture flat bottom well plates. After incubation for 48 h at 37 °C in a CO_2_ incubator, 50 µL of each cell supernatant were transferred into separate plates and added 50 µL of quantiblue reagent™ following manufacturer’s instructions (InvivoGen Inc.). Afterwards, this mixture was incubated for 3 h at 37 °C and then absorbance was measured at 650 nm. Results are expressed as absorbance values read at 650 nm in a plate reader (mutiscan Labsystems).

### Transmision Electron Microscopy (TEM) of promastigotes and bone marrow derived macrophages (BMM) infected with regular or DsRed *L. mexicana* and their treatment with urequinona

Promastigotes and BMM were treated with urequinona as described previously and were suspended in a solution of 3% glutaraldehyde in order to be processed for TEM (10). Both promastigotes and BMM were individually spun down and washed twice with a buffer solution of sodium cacodylate (pH=7.4). Samples were further incubated for 1 h at room temperature with osmium tetroxide in sym-collidine buffer solution (pH=7.6). At this point samples were washed twice with sym coolidine buffer solution. Samples (promastigotes and infected BMM) pellets were treated with a saturated solution of uranyl acetate in water. Samples were then dehydrated with a serial treatment with ethanol at different dilutions (ranging from 0-100% ethanol). Next, sample suspensions were treated overnight with a spurr’s resin diluted in acetone (1:1 dilution, Electron Microscopy Sciences, Fort Washington PA). Samples were spun down and pellets were included in polyethylene capsules (BEEM,^™^ Merck) that contained spurr’s resin. Polymerization of these epoxy blocks was performed at 70 °C and then they were sectioned with an untramicrotome (Leica microsistems Gmbh, Wein Austria). Very small small sections of 80 nm thickness were collected on a cupper grid of 200 mesh (pore size, Electron Microscopy Sciences) and stained with lead citrate for 3 min and evaluated for utrastructural analysis. Regarding BMM infection with transgenic parasites macrophage suspension was maintained as previously described and was infected overnight with DsRed *L. mexicana* and then treated for 48 h with 10 µM of urequinona. A control culture included cells infected with transgenic parasites and treated with 2% of DMSO in complete media. Next, cells from control and experimental culture were washed with PBS and mounted with DAPI (4’,6-diamidino-2-phenylindole) over glass slides using slide mounting media from Sigma-Aldrich and observed under the fluorescent microscope. Intracellular killing of parasites can be assessed by fluorescence fading by observation under fluorescence microscope and flow sight testing as described below.

### Quantitation of cytokines in supernatants of splenocytes from infected mice or BMM after treatment with urequinona

We determined the concentration of IFN-γ and IL-2 in supernatants of splenocytes (pulsed *in vitro* with parasite antigen) from infected and urequinona treated mice using a capture ELISA assay. We also used this assay to measure the concentration of IL-6 and IL-12p70 in supernatants from infected BMM also treated *in vitro* with urequinona. ELISA assay was performed using different concentrations of recombinant cytokines as standards to plot a standard curve. We used specific capture and detection monoclonal antibodies (MoAb) from specific hybridoma clones from commercial sources (BDPharmingen, San Diego CA). Results were extrapolated from a standard curve after reading in an ELISA reader at 405 nm (12).

### Image based cytometry (flow sight analysis)

Fluorescence of transgenic parasites (DsRed *L. mexicana*) infecting BMM was recorded with the ImageStreamX; Amnis Corporation, Seattle, WA (image based cytometry by measuring transgenic promastigote fluorescence fading inside of phagocytes treated with urequinona or 2% DMSO in complete media. Cells were treated with this alkaloid for 48 h and then counter-stained with DAPI for nucleus staining and contrast. Images were acquired utilizing the 488 nm laser and 20X magnification; cell images were represented as bright field and red fluorescence. Side scatter (SSC) was collected in channel 3 (camera 1) at 560-595 nm filter. Doublets and larger cell clusters were eliminated from the analysis following manufacturer instructions. Data analysis was performed using IDEAS software (12)

### Statistical analysis

Statistics for data handling included the use of Mann-Whitney and Dunnet tests and the IC_50_ values were calculated using a LdP Line®.

## Results

### Molecular structure of 24-Methylcholesta-4-24(28)-dien-3-one (urequinona)

Figure 1 shows the molecular structure and IUPAC (International Union of Pure and Applied Chemistry) name of tested compound (24-methylcholesta-4-24(28)-dien-3-one). For simplicity in this work, we use the trivial name of urequinona to refer to this known sterol.

### Promastigote and amastigote killing after treatment with urequinona

In order to demonstrate urequinona effectiveness to kill *L. mexicana in vitro*, we grew promastigotes and amastigotes (inside of BMM) in complete culture media and treated them with different concentrations (0-100 µM) of this steroid or sodium stibogluconate (reference drug 0-400 µM) as we described it in material and methods. Table I describes the IC50 values of these compounds against promastigotes and amastigotes treated *in vitro*. Promastigotes treated with urequinona displayed significantly lower (p<0.05) values of IC50 compared to reference drug (Table I). Regarding amastigote killing as they were treated with urequinona, we found that infected BMM presented considerably killing of intracellular parasites. The IC50 of amastigotes treated with urequinona was also significantly lower (p<0.05) compared to reference drug (Table I). Furthermore selectivity index was significantly higher (p<0.05) in cells treated with 10 µM of urequinona compared to reference drug (Table I). In a separate experiment we showed that the average number of amastigotes (infection rate) and percentage of infected BMM (infection level) in control cultures was significantly (p<0.05) higher compared to infected macrophages treated with 10 µM of urequinona for 48 h (Supplementary Fig 1).

### Transmission Electron Microscopy (TEM) of promastigotes and amastigotes *L. mexicana* and evaluation of loss of metabolic activity in DsRed *L. mexicana* amastigotes treated with urequinona

To evaluate ultrastructural damage of promastigotes and amastigotes as well as loss of metabolic activity of the latter, we undertook the following set of experiments. Figure 2 shows transmission electron microscopy (TEM) images of a control representative promastigote only treated with 2% DMSO in complete media (Fig 2A) and a representative promastigote treated with 10 µM of urequinona for 48 h (Fig 2B). Figure 2C shows a TEM image of a representative control BMM only treated with 2% DMSO in complete culture media and Fig 2D displays TEM image of a representative BMM fed with parasites and then treated with 10 µM of urequinona for 48 h. It can be seen a lower number of amastigotes inside of parasitophorous vacuoles in infected BMM treated with urequinona as compared to the control BMM image. Figure 3 shows control DsRed *L. mexicana* ingested by BBM only treated with 2% of DMSO in complete media (Fig 3A,C). Interestingly, transgenic parasites ingested by BMM showed fading of fluorescence after treatment with 10 µM of urequinona for 48 h (Fig 3B,D). This is an indication of parasite killing due to loss of metabolic activity since red fluorescent protein in transgenic parasites stops its production and glowing when parasites die. BMM nuclei (blue color stained) were counter-stained with DAPI for contrast.

**Fig 2.**
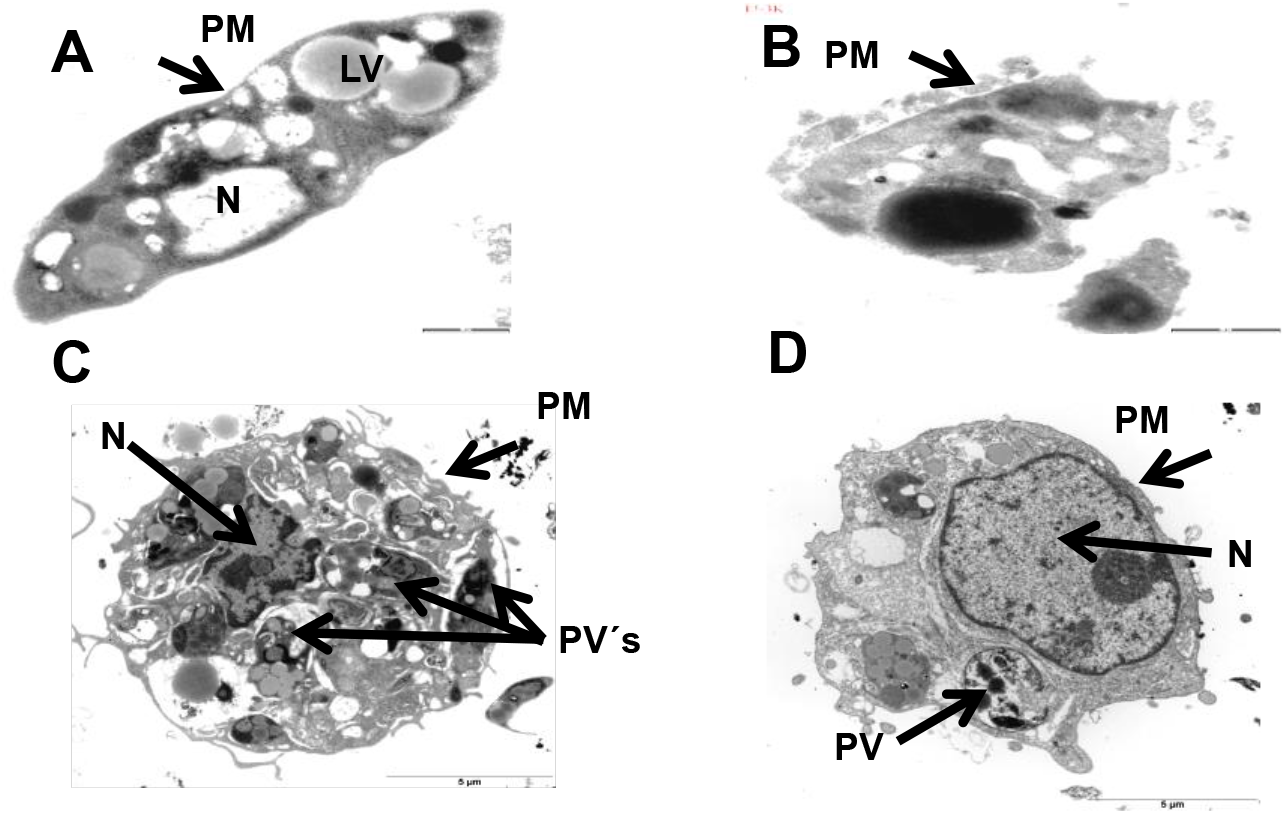
Ultraestructure of promastigotes and amastigotes inside of BMM. Control of representative promastigote and infected macrophage treated with 2% DMSO (A,C), representative promastigote and infected macrophage treated with 10 µM of urequinona for 48 h (B,D). Magnification bar of A and B is 1 µm whereas in C and D correspond to 5 µm. PM=Plasma membrane, N=Nucleous, PV=Parasitophorous vacuole, LV=Lipid vesicles.

**Fig 3.**
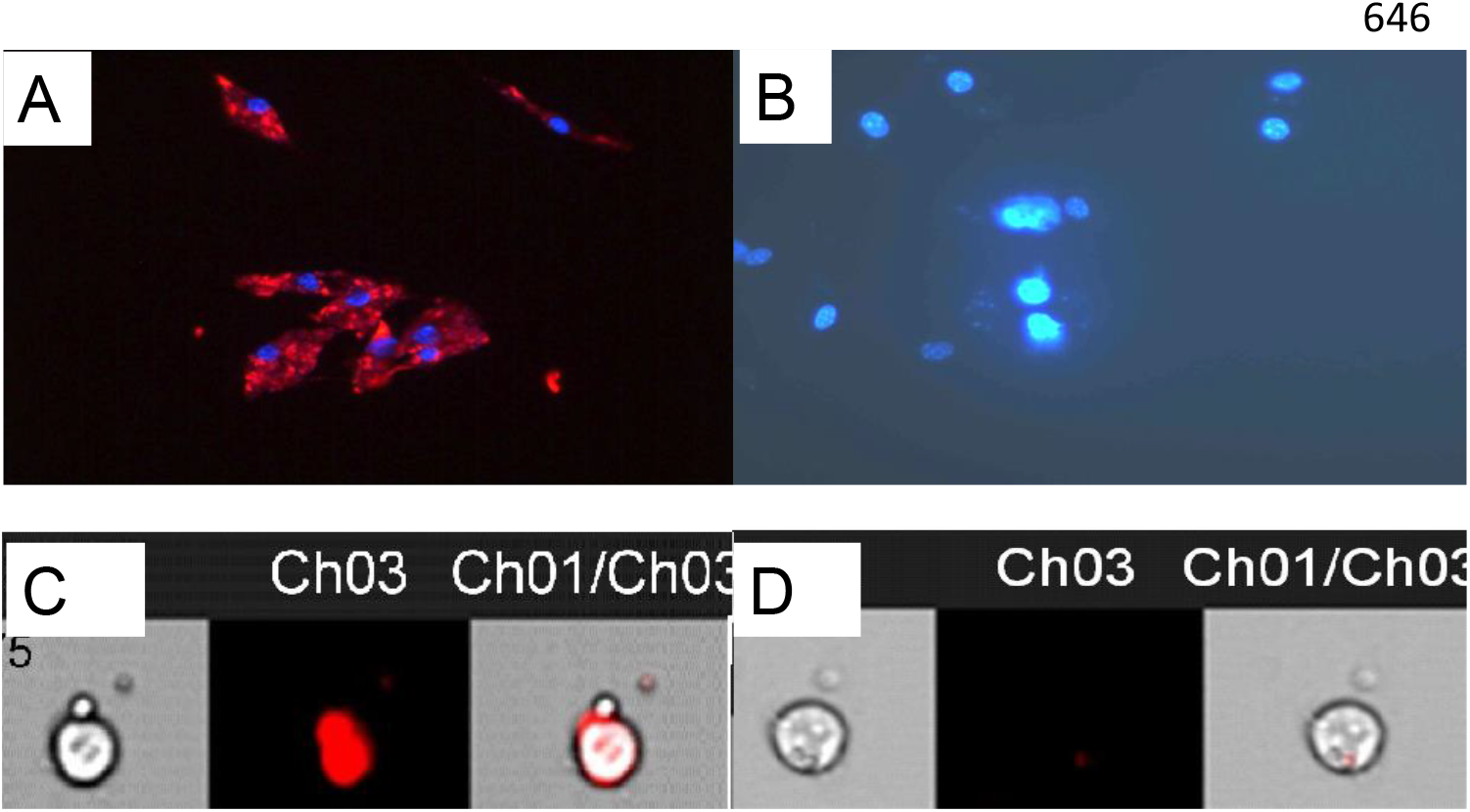
Amastigotes of DsRed*-L. mexicana* inside of BMM treated for 48 h with 2% DMSO (A, C) or 10 µM of urequinona (B, D). BMM s nucleous of Figs A and B were stained with DAPI for contrast and appear in blue. A and B images taken with a fuorescence microscope (100X). Figure also shows bright transgenic parasites (C) that fainted after treatment with urequinona (D). Images (C and D) were taken with a ImageStreamX.

### Programed cell death (apoptosis) and necrosis of promastigotes treated with urequinona

In order to find out if parasites treated with urequinona undertake a necrotic or an apoptotic pathway we designed the following experiments: Promastigotes of *L. mexicana* were treated with 10 µM of urequinona for 48 h as described in material and methods. Results show that parasites cultured for 48 h, once they reached the stationary phase of growth, presented apoptosis and necrosis death due to staining with anexin V and propidium iodide respectively (Fig 4B).

**Fig 4.**
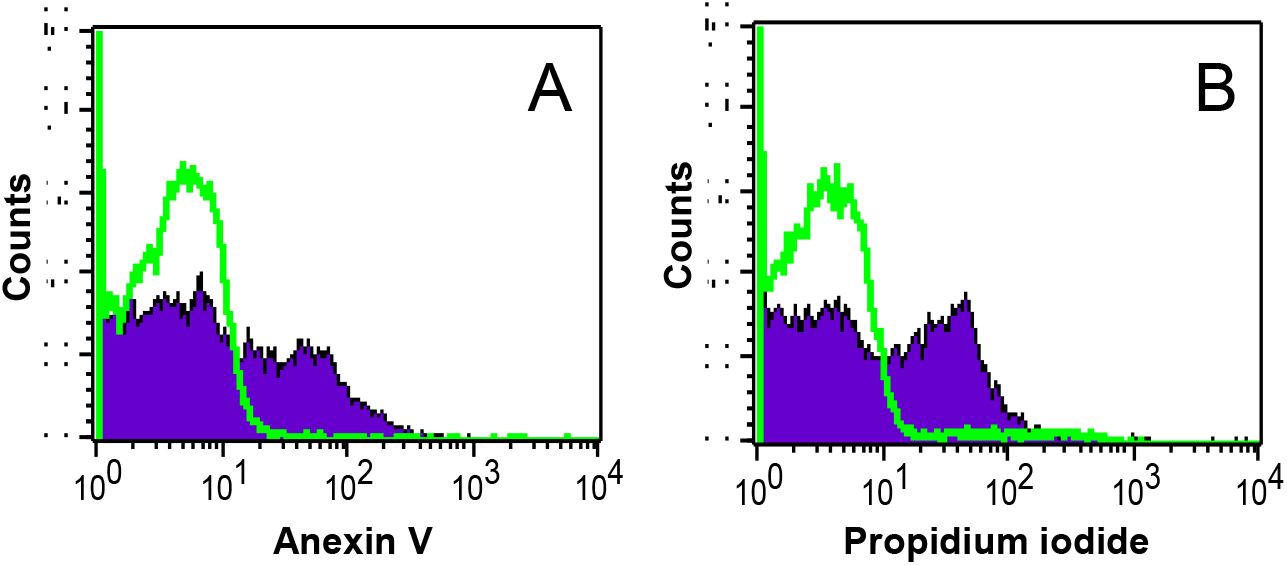
Apoptosis (A) and necrosis (B) of promastigotes. Open histogram shows control samples treated with 2%DMSO while close histogram shows parasites treated with 10 µm of urequinona for 48 h and stained with anexin V (A) or propidium iodide (B).

### Synthesis of IFN-γ and IL-2 by splenocytes of infected and urequinona treated mice

In figure 5 we show levels of IL-2 and IFN-**γ** produced by splenocytes from infected mice treated or not with topical preparation of urequinona and stimulated with parasite antigen *in vitro*. It can be observed that splenocytes from infected mice treated *in vitro* with 1 µg of protein from parasite antigen produced significantly (p<0.05) more IL-2 and IFN-**γ** compared to splenocytes from control mice only treated with vehicle and stimulated *in vitro* with 1 µg of protein from parasite antigen. This figure also shows control splenocytes treated with the mitogen Con-A as positive indicator of cell proliferation (Fig 5).

**Fig 5.**
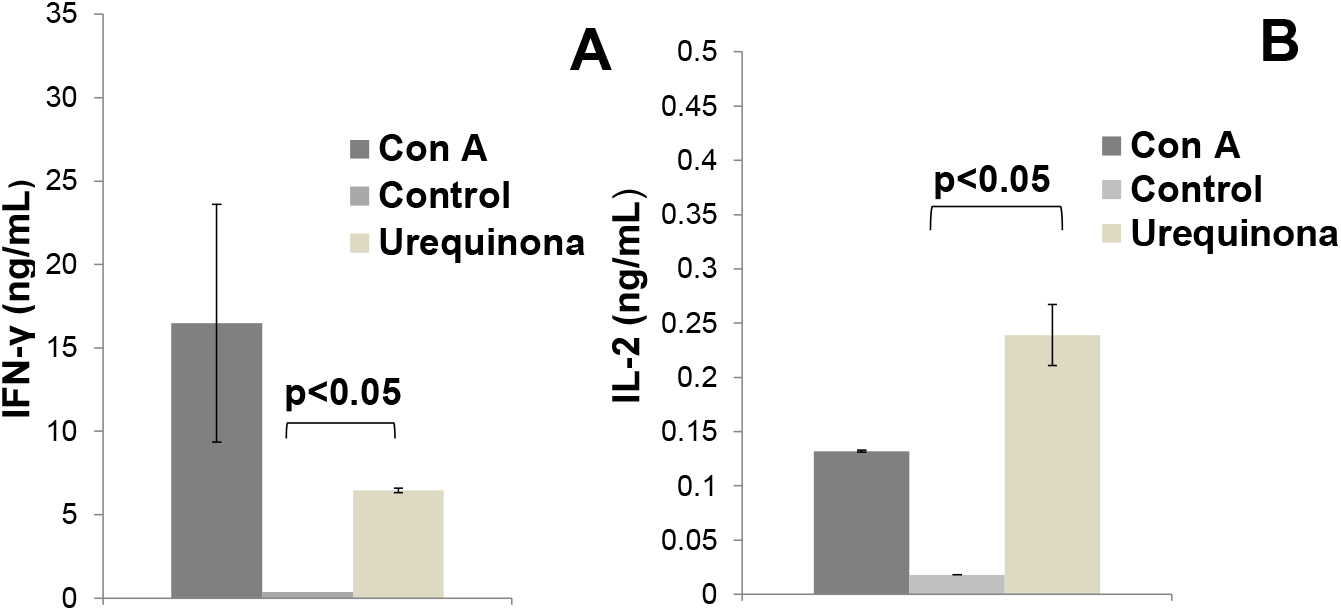
Concentration of IFN-γ (A) and IL-2 (B) produced by splenocytes from infected and urequinona treated mice and followed by *in vitro* stimulation with 1 µg of parasite antigen. Control group was treated with 2% DMSO and Con A treated group was included as a control of proliferation n=4.

### Activation of NFkB/AP-1 transcription factors and production IL-6 and IL-12p70 of infected BMM treated with urequinona

We were also interested to find out activation of the transcription factor NFkB/AP-1 in macrophages and the production of pro-inflammatory IL-6 and IL-12p70 after treatment with urequinona for 48 h. To this end we treated un-infected RAW reporter macrophages with 10 µM of urequinona or 2% of DMSO included as control. Supernatants of urequinona treated cells presented significant enhancement (p<0.05) of activation of this transcription factor compared to control cells. We also found a significant reduction (p<0.05) of the concentration of IL-6 (Fig 6B) and IL-12p70 (Fig 6C) in un-infected Raw macrophages treated with 10 µM of urequinona for 48 h compared to control cultures.

**Fig 6.**
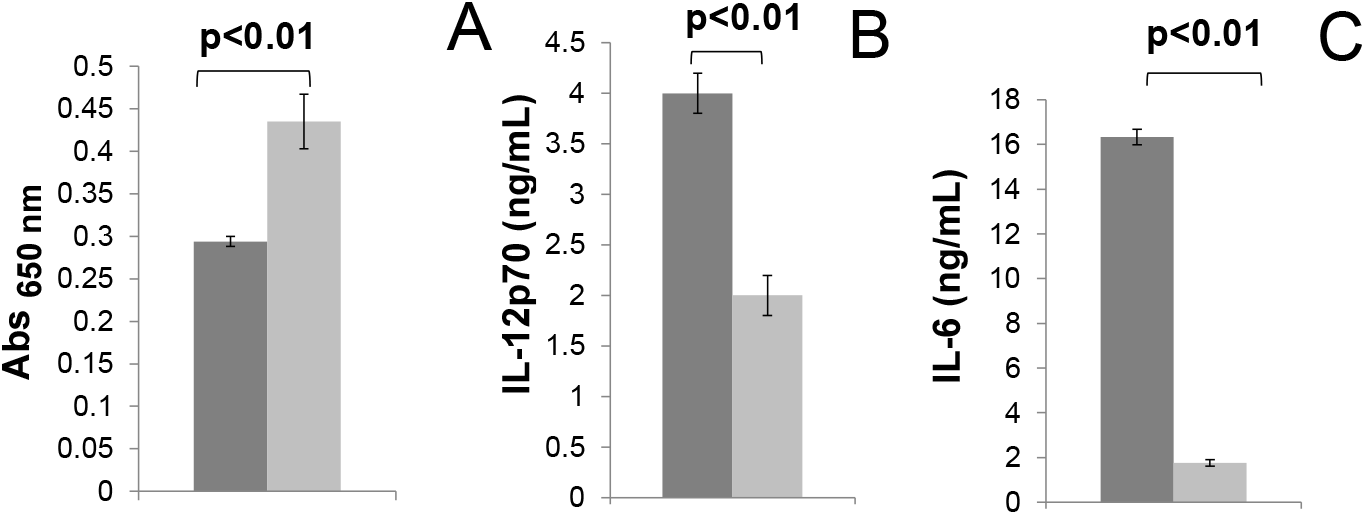
Activation of transcription factors NFkB/AP-1 (A) and production of IL-12p70 (B) and IL-6 (C) by infected BMM. Dark grey bars correspond to supernatants from control BMM treated with 2% DMSO and light grey bars correspond to supernatants from BMM treated with 10 µM of urequinona, n=4.

### Treatment of *L. mexicana* infected ears of C57BL/6 mice with the topical formulation of urequinona

To find out urequinona effectiveness to heal mice treated with low doses of topical preparation of this steroid, we infected the right ear of animals and once lesions were visible and palpable we daily treated them with 10 µM of urequinona formulation for 12 weeks. Results displayed in Fig 7A show that animals treated with the formulation of urequinona presented a significant reduction in lesion size throughout the treatment period. This ear model of infection also showed significant (p<0.05) reduction of parasite number in ear’s tissue of animals treated with topical formulation of urequinona compared to control animals only treated with vehicle solution (Figure 7B). This figure also shows histological preparations of a randomly chosen infected and vehicle-treated mouse ear showing abundant parasites (Fig 7C) and the one treated with the urequinona formulation display very few parasites (7D).

**Fig 7.**
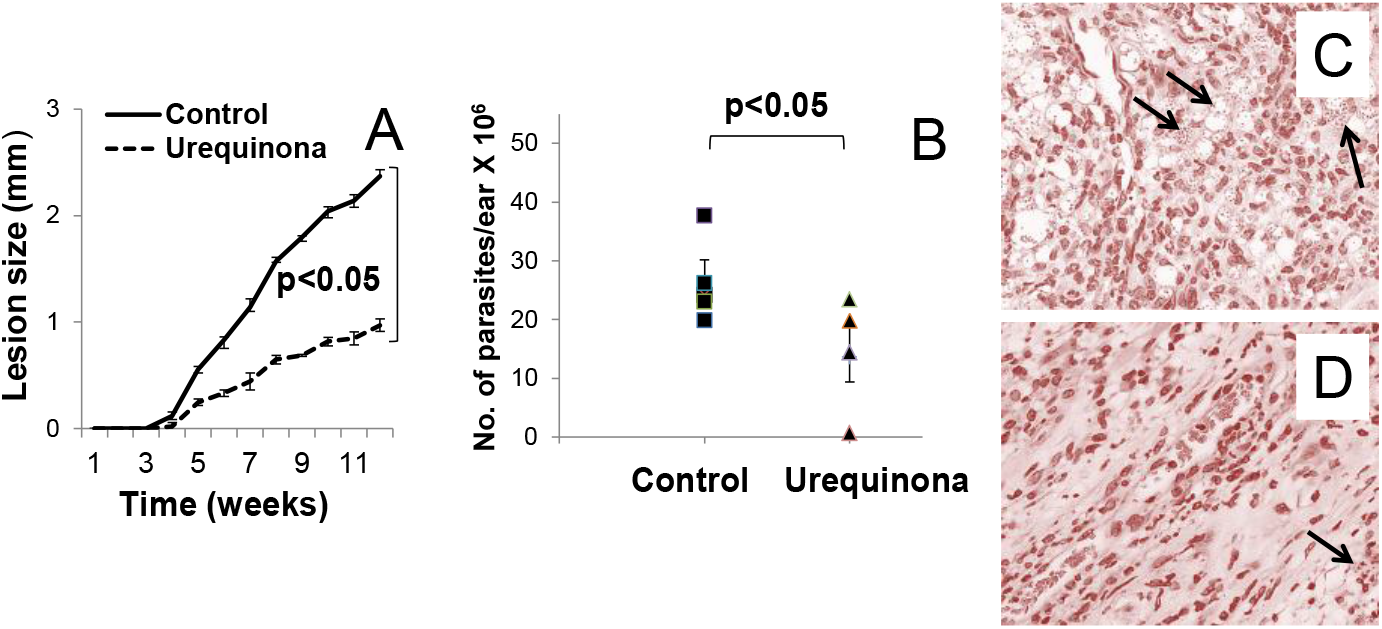
Lesion growth in C57BL/6 mice. Continuos line shows untreated animals while dotted line shows animals daily treated with 10 µM of urequinona/ear (n=4, A). The number of parasites recovered from individual ears were counted (B). Histology of a representative ear from a control mice is shown (C) and one treated daily for 12 weeks with 10 µM of urequinona/ear is also shown (D). Arrows show abundant parasites in control tissue sample while few parasites are present in urequinone treated tissue sample.

**Table 1.**
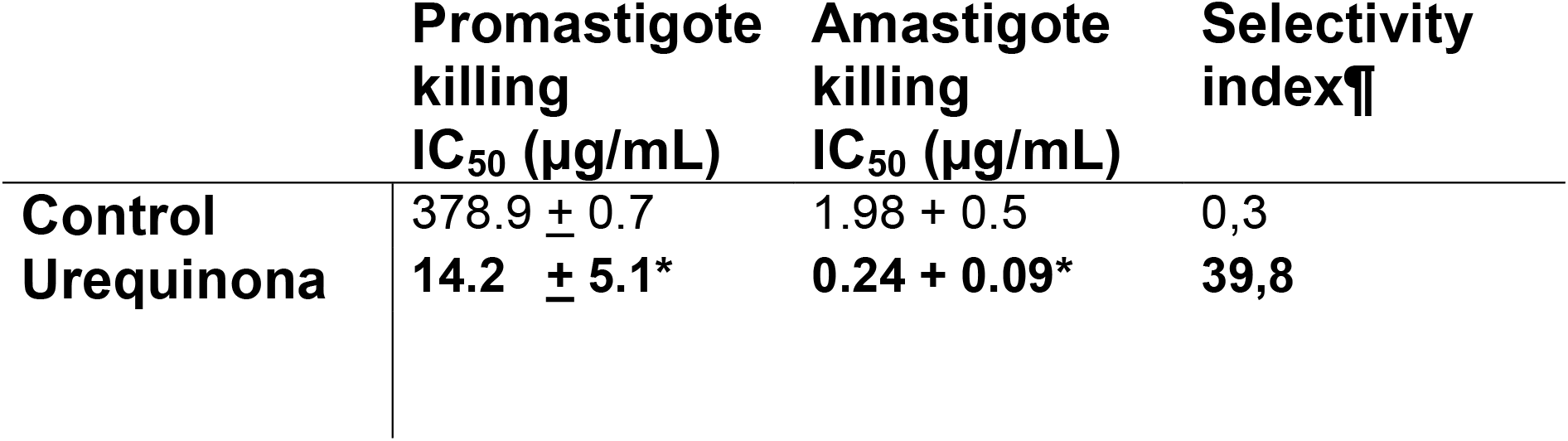
IC_50_ of promastigotes and amastigotes treated with urequinona. ¶Selectivity index is the ratio between cytotoxicity (50% of killing) of urequinona against uninfected BMM and average number of amastigotes in BMM. IC_50_ is the amount of urequinona that kills 50% of promastgotes or amastigotes *(p<0.05 compared to control). As reference drug we use sodium stibogluconate (control).

## Discussion

In Mexico antimonial drugs constitute the first line of treatment against LCL and when available patient’s treatment relies in Mexican government distribution. They are not available in drug stores and they present increasing levels of toxicity and parasite resistance (11-13). These limitations clearly highlight the need of affordable and readily available new drug treatments for LCL. In this work we undertook a series of experiments aimed to find out effectiveness of urequinona in hydroxyethyl cellulose as a topical alternative to treat LCL in C57BL/6 mice. We also studied several pathways induced by this molecule that contribute to control this disease in infected C57BL/6 mice. Urequinona induce killing of promastigotes of *L. mexicana* following two different mechanisms: necrosis and apoptosis. Electron microscopy images show extensive damage to promastigote membrane structure. Amastigotes presented lower numbers in parasitophorous vacuoles and irregular morphology as that is an indication of intracellular parasite death. We also observed parasite fluorescence fading in BMM infected with transgenic parasites treated with urequinona that indicates parasite’s loss of metabolic activity and death. Spleens from infected mice treated with urequinona synthetized significantly higher levels of IL-2 and IFN-γ, while BMM infected *in vitro* and then treated with urequinona presented lower levels of IL-6 and IL-12p70 compared to controls. Activation of transcription factors NFkB/AP-1 was enhanced in reporter RAW macrophages treated with urequinona. It is interesting that all cytokines studied in this work are pro-inflammatory molecules but they are differentially synthetized in lymphocytes and macrophages perhaps due to different parasite behavior inside of phagocytes compared to lymphocytes where parasites are not internalized. We can speculate that urequinona diffuses inside of parasitophorous vacuoles full of parasites and alter the balance between pro- and anti-inflammatory cytokines leading to efficient intracellular parasite killing. NFkB/AP-1 transcription factors are involved in expression of genes that code for control of cellular growth, cellular apoptosis, inflammation and lymphoid differentiation (14-16). Our data also show that urequinona induces production of IL-2 and IFN-γ in splenocytes from infected mice. IL-2 is a T cell growth factor that is essentially needed for T cell growth and differentiation of memory and effector cells (17-19). IFN-γ is a critical factor inducing a number of host protective responses such as enhancing antigen presentation and processing, it increases leukocyte trafficking, cellular proliferation, apoptosis and protection against infectious agents (20,21). Therefore by inducing production of these two important molecules we can assume urequinona up regulates the anti-parasite immune response. Interestingly our data show that urequinona decreases production of pro-inflammatory cytokines such as IL-6 and IL-12p70 in *in vitro* infected BMM. IL-12p70 and IL-6 are pleotropic cytokines with pro-inflammatory properties. It is well documented that IL-12p70 induces intracellular killing of different species of intracellular *Leishmania* parasites while IL-6 activates T lymphocytes among many other immunological functions (3,4,18,23,24). Human work we performed in the past demonstrated that IL-12p40, IL-2 and TNF-α plasma concentration increase while IL-6 is reduced in humans with active *L. mexicana* infections while IL-12p70 plasma levels remained unchanged (3,4). Therefore is not surprising that IL-6 and IL-12p70 levels are reduced and IL-2 is increased in urequinona stimulated infected BMM. Perhaps differential regulation of cytokines produced by various cell lineages could account for this observation. These findings would not be surprising since *P. andrieuxii* phytosteroids present biological differences to human steroids when they stimulate promastigotes or amastigotes *in vitro* (11). For example urequinona is a steroid with killing properties over *L. mexicana* parasites (10). In contrast with human steroids such as estradiol (E2), testosterone (T) and dihydrotestosterone (DHT) that do not kill these parasites but induce macrophages to destroy them (E2) or exacerbate (T and DHT) their intracellular reproduction (25). We believe urequinona encapsulated in hydroxyethyl cellulose for topical treatment of LCL would be a novel alternative to toxic licensed drugs currently in use to treat LCL by parenteral administration.

Authors declare there is not conflict of interest.

## Acknowledgements

Authors express their gratitude to Ohio State University for their facilities provided to perform this project. Dr Lezama-Dávila worked at OSU as postdoctoral fellow from 2005-2007 and Research Scientist from 2009-2013.

---

**Supplementary Fig1.**
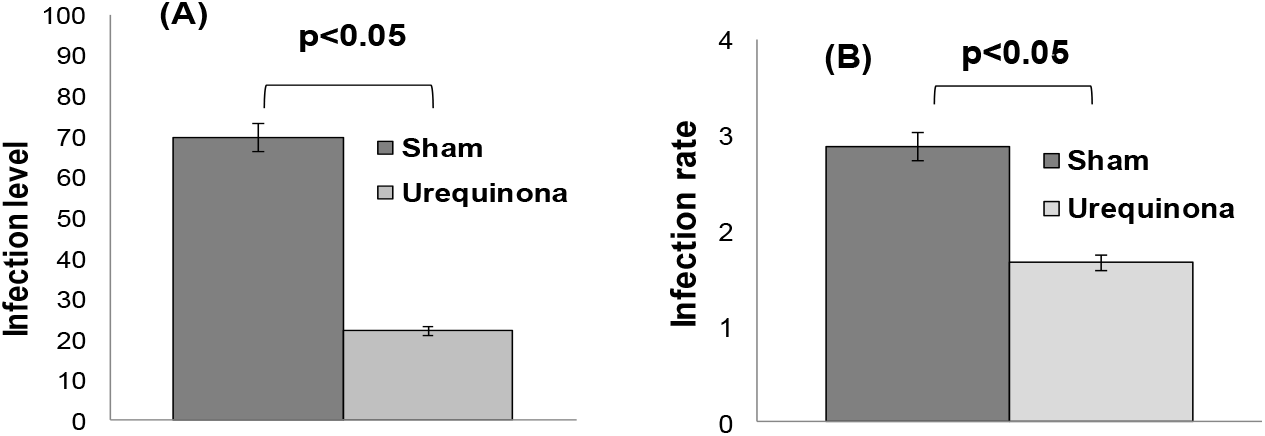
Bone marrow macrophages (BMM) infection with *L. mexicana* followed by daily treatment for 12 weeks with 10 µM of urequinona. Infection level is the percentage of infected BMM (A) and infectious rate is the average number of amastigotes per BMM (B)

